# Epigenetic reprogramming restores metronidazole sensitivity in drug-resistant *Trichomonas vaginalis* parasites

**DOI:** 10.1101/2025.01.07.631743

**Authors:** Julieta Seifert-Gorzycki, Daniela Muñoz, Loic Ciampossin, Ayelen Lizarraga, Suhani Bhanvadia, Lucrecia Iriarte, Samira Elikaee, Verónica Coceres, Patricia J. Johnson, Karine G. Le Roch, Pablo H. Strobl-Mazzulla, Natalia de Miguel

**Author notes:** For correspondence (NdM). These authors contributed equally to this work (Julieta Seifert-Gorzycki, Daniela Muñoz).

## Abstract

Resistance to antimicrobial drugs is a major global health challenge, yet the regulatory mechanisms underlying resistance phenotypes remain incompletely understood. In particular, whether epigenetic processes contribute to antimicrobial resistance in microbial eukaryotes is largely unexplored. Here, we show that epigenetic regulation plays a central role in modulating metronidazole resistance in *Trichomonas vaginalis*, the causative agent of the most common non-viral sexually transmitted infection. Pharmacological inhibition of histone deacetylases restores drug sensitivity in a highly resistant clinical isolate, markedly reducing the minimum inhibitory concentration. Integrative transcriptomic reveal widespread transcriptional reprogramming associated with activation of pathways involved in redox metabolism, transcriptional regulation, and stress responses. Chromatin profiling demonstrates that increased H3K9 acetylation is strongly associated with transcriptional activation. Upon TSA treatment, increased H3K9 acetylation is observed at promoters and gene bodies of genes that may be associated with MTZ resistance. By combining epigenetic perturbation with comparative transcriptomics across multiple strains, we identify a set of genes that are epigenetically silenced in resistant parasites that are constitutively expressed in sensitive backgrounds. Functional re-expression of two candidates, thioredoxin reductase and a Myb-like transcription factor (Myb6), significantly reduces resistance, establishing a direct link between epigenetic regulation and drug sensitivity. Together, our findings uncover a reversible, chromatin-based mechanism underlying antimicrobial resistance in a protozoan parasite and highlight epigenetic reprogramming as a potential therapeutic strategy. More broadly, this work suggests that epigenetic regulation may represent a general and previously underappreciated layer of control over drug resistance in microbial eukaryotes.

## INTRODUCTION

Eukaryotic microbial pathogens are major contributors to global morbidity and mortality. While drug therapies have proven effective against many infections, the emergence of drug resistance underscores the urgent need for alternative treatments. *Trichomonas vaginalis* is the causative agent of trichomoniasis, the most prevalent non-viral sexually transmitted infection worldwide, with an estimated 156 million new cases annually ^1^. Infection is associated with vaginitis, adverse pregnancy outcomes ^2^, pelvic inflammatory disease, and has been linked to increased risk of malignant cervical and prostate cancers ^3–5^. Despite its substantial global burden, treatment options for trichomoniasis are extremely limited.

The 5-nitroimidazole drugs, particularly metronidazole (MTZ) and tinidazole, are the only approved treatments for trichomoniasis with MTZ being the most widely prescribed ^6^. Resistance of *T. vaginalis* to MTZ, first reported in 1962 ^7^, has been increasingly documented ^8^, particularly in persistent or recurrent infection, highlighting the urgent need to better understand the mechanisms driving resistance. MTZ functions as a prodrug, requiring reduction of its nitro group to exert its antimicrobial effects ^8^. It damages DNA, forms protein adducts and induces oxidative stress by depleting thiol pools ^9^. Resistance mechanisms fall into two categories: aerobic resistance, associated with oxygen stress responses, and anaerobic resistance, linked to hydrogenosome metabolism deficiencies ^10^. Aerobic resistance involves enhanced activity of oxygen-scavenging enzymes (e.g., superoxide dismutase, flavoprotein oxidases), which protect the parasite from oxidative stress. In this context, oxygen serves as the preferred electron acceptor, preventing MTZ reduction and activation in hydrogenosomes ^9,11^. Anaerobic resistance, on the other hand, arises from downregulation of hydrogenosome enzymes such as pyruvate:ferredoxin oxidoreductase (PFOR) and ferredoxin (Fd) ^11–13^, critical for MTZ activation. Interestingly, certain proteins, such as flavin reductase 1 (FR1), are implicated in both resistance types, as they are often absent or downregulated in both aerobic and anaerobic metronidazole-resistant isolates ^9,12^. MTZ resistance studies have primarily focused on candidate genes related to oxygen scavenging and hydrogenosome metabolism ^12^. Despite significant progress, disruptions in known genes associated with MTZ resistance do not fully explain this phenomenon in all cases, suggesting that it is a multifactorial process. In this sense, large-scale genetic analyses revealed specific single nucleotide polymorphisms ^14,15^ and strain-specific gene expression differences in metabolic pathways associated with resistance have also been implicated ^15–17^. Notably, resistant parasites often display broad transcriptional changes rather than discrete gene mutations, suggesting that regulatory mechanisms beyond fixed genetic alterations may contribute to drug tolerance and resistance.

Epigenetic regulation provides a powerful and reversible means of controlling gene expression in response to environmental cues, including drug pressure. In other eukaryotic systems, most notably cancer cells and fungal pathogens, epigenetic mechanisms such as histone acetylation have been shown to drive transient drug tolerance and reversible resistance states ^18–20^. In cancer cells, subpopulations can adopt a reversible drug-tolerant phenotype maintained by altered chromatin states, and disruption of these states with epigenetic agents restores drug sensitivity ^21,22^. Inhibition of histone deacetylases (HDAC) promotes the accumulation of acetylated histones, leading to a more open chromatin configuration that facilitates transcriptional reactivation of previously silenced genes. HDAC inhibitors (HDACi) have therefore emerged as promising therapeutic agents for treating cancers, immunological diseases and other conditions as they can block angiogenesis, arrest cell growth and lead to differentiation and apoptosis in tumor cells ^23–25^. In contrast, despite their well-established effects on chromatin dynamics, the contribution of HDACs and histone acetylation to antimicrobial drug resistance in infectious diseases remains poorly understood. In *T. vaginalis,* previous findings demonstrated that histone acetylation influences both transcription and pathogenesis ^26–28^. Specifically, treating a poorly adherent strain of *T. vaginalis* with the HDACi Trichostatin A (TSA) increased H3KAc and chromatin accessibility around transcription start sites, upregulating genes involved in adherence ^26^. Although chromatin-based regulation is increasingly recognized as a major determinant of transcriptional control, whether epigenetic silencing or activation of resistance-associated pathways influences MTZ susceptibility in this parasite remains unknown. Here, we investigate the role of epigenetic regulation in modulating MTZ resistance in *T. vaginalis*. Using a highly resistant clinical isolate, we combine pharmacological inhibition of histone deacetylases with genome-wide transcriptomic and chromatin profiling to examine how histone acetylation shapes transcriptional responses to drug exposure. By integrating these data with comparative transcriptomics from multiple MTZ-sensitive strains and performing functional validation of candidate genes, we identify epigenetically regulated pathways that directly influence drug sensitivity. Our findings reveal epigenetic reprogramming as a key determinant of MTZ resistance and uncover a previously unrecognized, reversible layer of antimicrobial resistance control in a major human protozoan parasite.

## MATERIALS AND METHODS

### Parasites, cell cultures, and media

*Trichomonas vaginalis* strains BRIS/92/STDL/B7268 ^29,30^, SD7 ^31^, CDC1132 (MSA1132; Mt. Dora, Fla, USA 2008) ^32^ and NYH209 (ATCC 50146) ^33^ were cultured in Tryptose Yeast Maltose (TYM) medium ^34^ supplemented with 10% fetal bovine serum and 10 U/ml Penicillin/Streptomycin (Invitrogen). B7268 strain was obtained from a Brisbane patient with refractory infection that displays both high aerobic and anaerobic resistance to metronidazole ^30^. All parasite strains were grown at 37°C and passed daily.

### Metronidazole susceptibility assay and Trichostatin A treatment

Metronidazole (MTZ) susceptibility of *T. vaginalis* was determined by microscopy using a standard minimum inhibitory concentration (MIC) assay under microaerophilic conditions. To this end, log-phase parasites were seeded at 2 × 10^5^ parasites/ml and exposed to serial dilutions of metronidazole (0 - 400 µg/ml, Sigma-Aldrich) dissolved in DMSO. Final volumes were adjusted to ensure that the DMSO concentration did not exceed 0.1% (v/v), control tubes (0 µg/mL MTZ) received an equivalent volume of DMSO to match the maximum volume used in the treatment. Cultures were incubated for 48 h and examined by inverted light microscopy (40X) to assess parasite motility. The MIC was defined as the lowest MTZ concentration at which no motile parasites were observed. Loss of viability was confirmed by the inability of non-motile parasites to proliferate following reinoculation into drug-free medium. All assays were performed in four independent experiments.

For TSA treatment assays, B7268, CDC1132, and SD7 parasites were exposed to 100 nM trichostatin A (TSA, Sigma-Aldrich) for 16 h prior to MTZ treatment. After addition of MTZ, parasite growth was monitored for 48 h across increasing MTZ concentrations and compared with untreated controls.

### RNA-seq of T. vaginalis

*Trichomonas vaginalis* strain B7268 treated with TSA (100 nM), with metronidazole (MTZ; 5 µg/ml), or with a combination of TSA and MTZ, as well as untreated parasites as control were sequenced in triplicates. Total RNA was extracted from ∼5 × 10^6^ *T. vaginalis* according to the protocol outlined in the Total RNA Purification Kit (Norgen Biotek Corp.) following the manufacturer’s instructions. RNA quality and quantity were assessed using an Agilent Bioanalyzer with RNA integrity numbers (RINs) of >7 for all samples. The mRNA libraries were paired end (100 bp) sequenced with Illumina using TruSeq Stranded mRNA (Macrogen, Inc).

### Transcriptome analyses

After sequencing, 20-26 million reads were generated per RNA-seq library. For quality control of the paired-end sequencing data, the software FastQC ^35^ was used. The adapter sequence content was identified and trimmed using Trimmomatic ^36^. Then, HISAT2 ^37^ was used to align the RNA-seq data sets to the G3 2022 reference genome sequence ^38^, and the results showed an overall alignment rate of >90% for all libraries. In order to quantify the counts read for each gene, featureCounts ^39^ was used with default parameters. Principal components analysis was carried out to evaluate the variation through biological replicates. Using the DESeq2 package, it was possible to identify up- and down-log2 fold changes in gene expression ^40^. Differential gene expression and dispersion were examined using MAplot and plotDispEsts functions, respectively. Only changes where |log2FC| > 1 and padj < 0.05 were considered significant. Using the results command from DESeq2, the expression patterns of the three treatments (MTZ, TSA, and TSA_MTZ) were compared pairwise against the control group. In order to identify the genes differentially expressed in a susceptible strain to MTZ, RNA-seq dataset from wild type sensitive (NYH209) and resistant (B7268) strains were obtained from Sequence Read Archive (SRA) in online repositories (BioProject accession code PRJNA1188544). For the final list of candidates genes a Gene Ontology (GO) enrichment analysis was carried out using the TrichDB database of open access ^41^. P-value ≤ 0.05 was considered significantly enriched. Additionally, publicly available RNA-seq datasets from nine sensitive *T. vaginalis* strains were downloaded from the Sequence Read Archive (accession codes SRP057357 and SRP057311) ^15^ and reanalyzed as described above to enable comparative analysis with the resistant strain B7268. The overall alignment rate to the G3 reference genome for all downloaded samples exceeded 90%.

### Parasite viability assays

Parasite viability was analyzed using the fluorescent exclusion dye propidium iodide (PI). For this purpose, wild-type B7268 parasites (control) and B7268 parasites treated with 100 nM TSA for 16 hours, 5 µg/mL MTZ, or a combination of TSA and MTZ were labeled with 20 µg/mL propidium iodide (PI) at 4°C for 10 minutes. PI fluorescence associated with non-viable cells using paraformaldehyde (PFA) 4% (control of dead cells) was measured by flow cytometry. Cells were excited with 480 nm light, and emission was detected using a 585/42 nm filter (PI fluorescence, FL2) on a FACSCalibur flow cytometer (Becton-Dickinson). Data were analyzed using Flowing software version 2.4.1 (Perttu Terho, Turku Centre for Biotechnology, Finland; www.flowingsoftware.com). Four independent experiments were performed.

### ChIP assay

The ChIP assay was performed as described ^42^ with minor modifications for *Trichomonas vaginalis.* Briefly, *T. vaginalis* B7268 parasites treated with TSA+MTZ (100 nM + 5 µg/mL), or untreated (control) were cross-linked with formaldehyde (1%) for 10 min at 37°C, quenched with glycine (125 mM), and washed with PBS. Nuclei were extracted in HEPES-based buffer and lysed by passage through a 26 G needle. Chromatin was sheared using a Covaris S220 Ultrasonicator (6 min; duty cycle 10%; peak power 140; cycles per burst 200; 6°C), diluted in ChIP dilution buffer, and centrifuged to recover soluble chromatin. After pre-clearing with protein A agarose/salmon sperm DNA beads, 10% of each sample was reserved as input. Immunoprecipitation was performed overnight at 4°C using 2 µg of anti-H3K9Ac antibody (Diagenode, C15410004). Immune complexes were captured with blocked protein A beads and washed sequentially with low-salt, high-salt, and TE buffers. Chromatin was eluted, cross-links were reversed overnight at 45°C, and samples were treated with RNase A and proteinase K. DNA was recovered by phenol–chloroform extraction and ethanol precipitation, purified with AMPure XP beads, and used for library preparation with the NEBNext Ultra II DNA Library Prep Kit (E7645L).

### ChIP-seq analysis

Quality control of the paired-end ChIP-seq data was performed using FastQC software. Adapter sequences and low-quality bases were removed using Cutadapt ^43^. The trimmed paired-end reads were aligned to the T. vaginalis G3 reference genome (TrichDB-68) using Bowtie2 in paired-end mode. Following alignment, SAM files were converted to BAM format and filtered using samtools _44_to retain only properly paired reads with mapping quality ≥10, removing unmapped and QC-failed reads. PCR duplicates were identified and marked using PicardTools MarkDuplicates to account for potential amplification bias. Normalized coverage tracks were generated using deepTools bamCoverage with counts per million (CPM) normalization and a bin size of 10 bp. Additionally, input-subtracted tracks were created using bamCompare to account for background signal and technical variation. All downstream ChIP signal quantifications were performed using input-subtracted CPM tracks (CPM - Input). Peak calling was performed using MACS2 in paired-end mode (BAMPE) with the input sample as control. A q-value threshold of 0.05 and fold-enrichment cutoff of 1.5 were applied, using a genome size of 180 Mb. Average profile plots were generated using deepTools ^45^ and ChIP-seq data was visualized using PygenomeTracks ^46^. Differential H3K9Ac enrichment between conditions was assessed using DiffBind _47_. A consensus peak set was defined by requiring peak overlap in at least two replicates per condition (minOverlap = 2), and read counts were quantified at consensus peaks using SummarizeOverlaps (bUseSummarizeOverlaps = TRUE). Library size normalization was performed using DESeq2-based size factors as implemented in DiffBind (dba.normalize). Sample-level reproducibility was evaluated by principal component analysis (PCA) and pairwise Pearson correlation matrix on the normalized count matrix. Differential binding analysis was performed in parallel using DESeq2 comparing Control versus MTZ-treated samples and Control versus TSA-treated samples.

To evaluate the genome-wide relationship between H3K9Ac occupancy and transcriptional activity, RNA-seq read counts were quantified across all annotated protein-coding genes using featureCounts and normalized using DESeq2 size-factor estimation. Per-condition mean normalized counts were computed across biological replicates. ChIP-seq peaks from each condition (consensus peak sets from MACS2) were assigned to genes by coordinate overlap using bedtools intersect; when a peak overlapped multiple genes, the gene with the greatest overlap length was retained. The mean H3K9Ac signal per peak was extracted from input-subtracted CPM bigWigs over peak intervals using bigWigAverageOverBed, and replicate scores were averaged prior to analysis. Genes with an assigned peak were ranked by their DESeq2-normalized mean expression and partitioned into ten equal-sized deciles (D1 = lowest expression, D10 = highest). For each decile, the mean ± standard error of the mean (SEM) of H3K9Ac scores was computed. The monotonic association between expression decile rank and mean ChIP score was quantified using the Spearman rank correlation coefficient. To validate that the observed correlation reflects a genuine association rather than an artifact of the decile grouping procedure, a random permutation control was performed: expression labels were randomly shuffled 20 times across genes, and the full decile analysis was repeated for each permutation. Each permuted result is shown as a light gray line in the figure, while the real data are displayed as a colored line with error bars. The mean ± standard deviation of Spearman r values across permutations is reported alongside the real correlation coefficient.

### Quantitative real-time polymerase chain reaction (qRT-PCR)

Parasites were exposed to 100 nM trichostatin A (TSA) for 16 h prior to total RNA extraction. Total RNA from ∼5x10^6^parasites was extracted using RNA extraction kit (Invitrogen) following the manufacturer’s instructions. Total RNA was treated with DNase I (Invitrogen) and reverse transcribed using SuperScript II reverse transcriptase and oligo (dT) primers (Invitrogen). Real-time PCRs were performed using Brilliant SYBR Green qPCR Master Mix (Roche), 450 nM concentration of each primer and 200-500 ng of cDNA in a 20 μl reaction volume using a StepOnePlus real-time PCR system. Parallel reactions performed without reverse transcriptase were included as negative controls. Amplification curve analysis was performed using the LinRegPCR software^45^. LinRegPCR determined the PCR efficiency from the exponential phase of individual amplification curves using an automated baseline subtraction algorithm independent of ground phase cycles. The efficiency-corrected initial target quantity (N0) was calculated for each sample using the common quantification threshold, mean PCR efficiency per assay, and individual Cq values. Gene expression ratios were calculated as N0_target/N0_Hmp24, where Hmp24 served as the housekeeping gene. For each sample, duplicate test reactions were analyzed for the expression of the gene of interest. The sequences of the primers used are: Thioredoxin-disulfide reductase protein (TVAGG3_0315040): Forward TGGAATACCTCACCCAAAGAAG Reverse GCCAGCCTGTGCAGTATAA; Protein of unknown function (TVAGG3_0172050): Forward CCACCTAATACCAGAGGATATTCATTA and Reverse CAGACATGTATCAGTTACCCTTGA; Nucleotide-diphospho-sugar transferases family (TVAGG3_1037610): Forward CCAGTACAACAGAATCACTCCAA and Reverse CGTAAACGTATGATGGCCAAATG; Glycosyl hydrolases family (TVAGG3_0371210): Forward TTCCTCATGGCGAAGGAATG and Reverse CTCGCTGCTGCATAGAATGA

### Plasmid construction and exogenous expression in *T. vaginalis*

Master-Neo-(HA) plasmids ^48^ containing the open reading frame of each candidate gene (TrxR and Myb6) were purchased from GeneScript (USA). Parasites (B7268 strain) were nucleofected with 15 μg of each construct or an empty vector control (EpNeo), as described previously ^49^ with the following modifications: parasites were spin down and washed with 5% sucrose (Sigma-Aldrich) in 1x PBS (pH 7.4) once before using SE Cell Line 4D-Nucleofector® kit (Lonza #V4XC-1032) and CM-150 program for nucleofection. After 10 min on ice, parasites were cultured in Diamond’s media for 24 hr. Transfectants were then selected and maintained using 100 μg/mL G418 (Sigma).

### Immunolocalization experiment

Parasites were incubated at 37°C on glass coverslips for 4□h. The parasites were fixed with PFA 4% and permeabilized in 0.1% Triton for 10□min. The cells were washed and blocked with 5% fetal bovine serum (FBS) in phosphate buffered saline (PBS) for 30 min, incubated with a 1:500 dilution of anti-HA mouse primary antibody (Covance, Emeryville, CA, USA) diluted in PBS 2%FBS, washed and then incubated with a 1:5000 dilution of Alexa Fluor-conjugated secondary antibody (Molecular Probes). The coverslips were mounted onto microscope slips using 4′, 6′-diamidino-2-phenylindole (Invitrogen). Fluorescence parasites were visualized using a Zeiss Axio Observer 7 (Zeiss) microscope.

### MYB Motif Enrichment and Positional Conservation Analysis

MYB-related motifs were downloaded from the JASPAR database ^50^ (total of 261 position weight matrices - PWMs). Motif enrichment analysis was performed using Analysis of Motif Enrichment (AME) from the MEME Suite ^51^. Target dataset comprised promoter sequences (defined as -60 to +40 bp relative to TSS) from 1,464 genes upregulated by both MTZ and TSA treatments (MTZ_TSA vs Control), while the control dataset consisted of promoter sequences from the remaining upregulated genes (974 genes from MTZ vs Control only, 477 genes from TSA vs Control only, and 1,893 genes from combined treatments; Fig. 3D). Statistical significance was assessed using Fisher’s exact test, and p-values were corrected for multiple testing using the Bonferroni method (261 motifs tested).

Positional conservation of MYB motif occurrences was examined using a custom Python implementation. The motifs underwent comprehensive positional scanning, where each promoter sequence was scanned using the motif PWM to identify all occurrences exceeding a likelihood ratio threshold. The genomic position (relative to TSS) of each motif match was recorded for both target and control groups. Motif occurrences were then grouped by their position relative to the TSS in 5 bp bins, and the frequency of genes containing motifs at each positional bin was calculated separately for target and control datasets. Positional groups containing motifs in more than 2 genes were retained for visualization. Positional enrichment was independently confirmed using CentriMo v5.5.9 ^52^ (Supplementary Figure 4).

## RESULTS

### Treatment with the histone deacetylase inhibitor TSA overcome metronidazole drug resistance in *T. vaginalis*

We propose that epigenetic modifications may silence genes involved in the development of metronidazole (MTZ) resistance. To test whether pharmacological modulation of chromatin structure can restore drug sensitivity, we treated a highly MTZ-resistant *T. vaginalis* strain (B7268) with the histone deacetylase (HDAC) inhibitor trichostatin A (TSA). MTZ susceptibility was then assessed using a standard minimum inhibitory concentration (MIC) assay under aerobic conditions. Resistant parasites were pretreated with 100 nM TSA for 16 hours, followed by MTZ treatment for 48 hours. This concentration of TSA has previously been shown to increase H3KAc, relax the chromatin and modulate gene expression in *T. vaginalis* ^26^. Our results show that TSA treatment markedly increased MTZ sensitivity in B7268-resistant parasites, reducing the MIC from 250 µg/ml to 8.75 µg/mL (p-value = 0.0002) (Fig. 1A). This sequential treatment with an epigenetic drug sensitized refractory B7268 parasites. To further validate these findings, we assessed the effect of TSA and MTZ in additional *T. vaginalis* strains, CDC1132 and SD7 (Fig. 1B). Although the effect was more pronounced in the highly resistant strain B7268, TSA treatment also increased drug sensitivity in CDC1132 and SD7 strains. In both strains, the MIC was decreased from 7.5 µg/ml to 3.125 µg/ml (p-value = 0.0069 and p-value=0.0143, respectively). Collectively, these finding suggest that TSA, possibly through histone acetylation and changes in chromatin accessibility, contributes to MTZ sensitivity in *T. vaginalis*.

**Figure 1:**
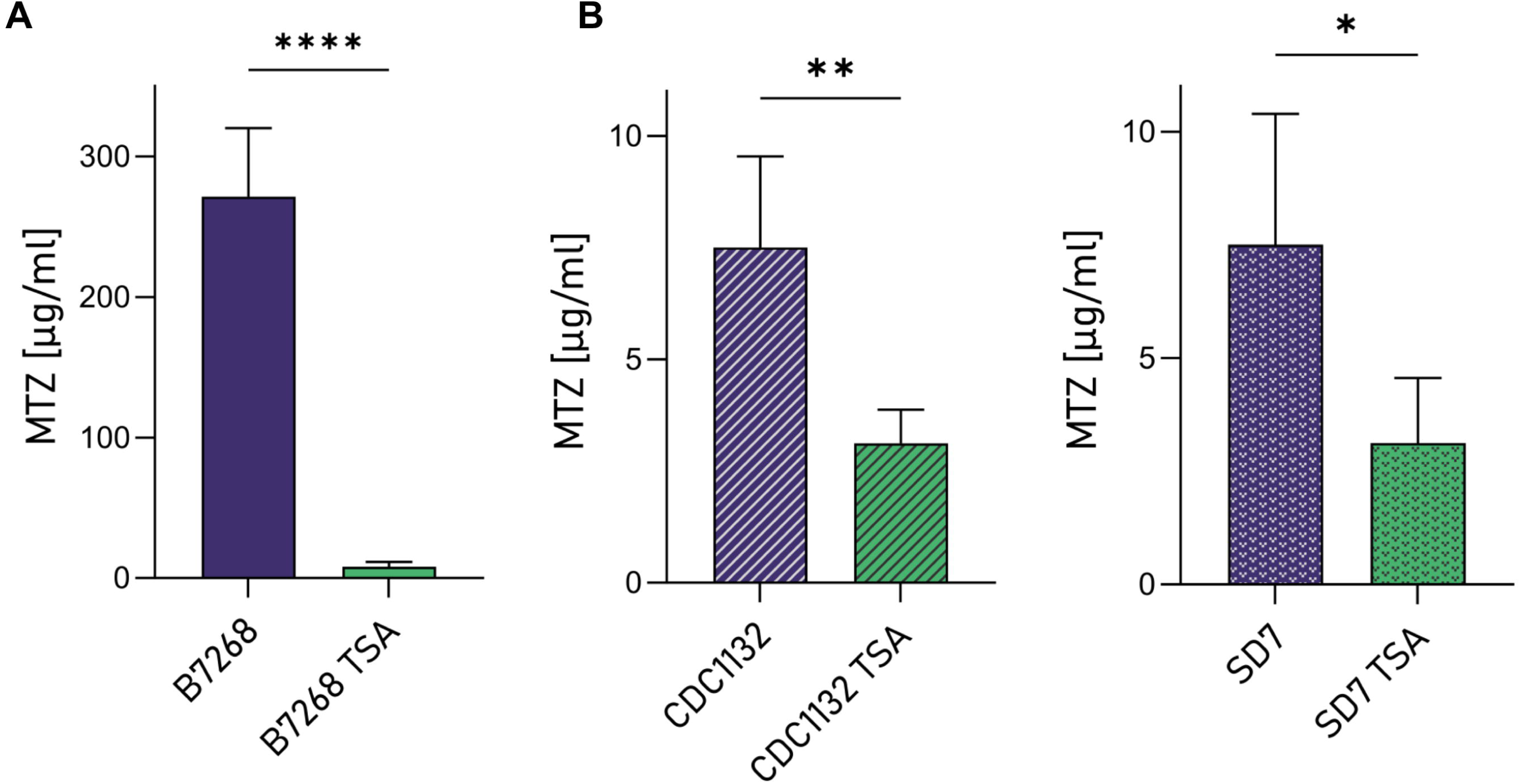
A. B7268 sensibilization to MTZ upon TSA pre-treatment. B. The sensibilization to MTZ upon TSA pre-treatment was also observed in CDC1132 and SD7 *T. vaginalis* strains. For each strain four independent experiments were conducted. Data are expressed as concentration of MTZ [μg/ml] for which the parasites were no longer motile ± standard deviation of the mean. Student T-tests (α=0.95) were used to determine significant differences between treatments. (Signif. codes: ns = P-value > 0.05, * = P-value ≤ 0.05, ** = P-value ≤ 0.01, *** = P-value ≤ 0.001, **** = P-value ≤ 0.0001)

### RNA-seq uncovers histone acetylation dependent pathways associated with MTZ resistance

Histone acetylation is a well-established epigenetic mechanism that promotes chromatin relaxation, thereby facilitating transcription factor access and enhancing gene expression. To examine the molecular basis underlying TSA-mediated modulation of MTZ resistance, we conducted whole-transcriptome RNA-seq analysis of the highly resistant *T. vaginalis* B7268 strain under four different conditions: untreated control, TSA (100 nM), MTZ (5 µg/ml), and sequential TSA followed by MTZ treatment (Fig. 2A). These conditions were selected as parasite viability remained unaffected across all treatments (Supplementary Fig. 1A). RNA-seq libraries were generated in biological triplicates. Principal component analysis (PCA) revealed clear separations among treatment groups, with PC1 and PC2 accounting for 59% and 20% variance, respectively, while demonstrating minimal variability among biological replicates (Supplementary Fig. 1B). Notably, transcriptomes from TSA-treated parasites clustered more closely with those from the sequential MTZ_TSA treatment than with MTZ-treated or untreated controls (Supplementary Fig. 1B). Consistent with this observation, the heatmap of differentially expressed genes revealed a distinct gene expression pattern between treated and untreated groups (Fig. 2B). To identify genes whose expression was modulated by TSA and potentially involved in MTZ resistance, we performed differential expression analysis comparing each treatment condition (MTZ, TSA and MTZ_TSA) with untreated B7268 controls and visualized the results using volcano plots (Fig. 3A-C and Supplementary Table 1). MTZ treatment alone resulted in 2,867 upregulated and 1,725 downregulated genes relative to control cells (Fig. 3A and Supplementary Table 1). TSA treatment led to a broader transcriptional activation, with 3,192 genes upregulated and 1,349 downregulated (Fig. 3B and Supplementary Table 1), consistent with HDAC inhibition predominantly promoting gene expression. Sequential TSA_MTZ treatment produced the most extensive transcriptional response, with 3,293 upregulated and 2,088 downregulated genes compared to untreated parasites (Fig. 3C and Supplementary Table 1).

**Figure 2:**
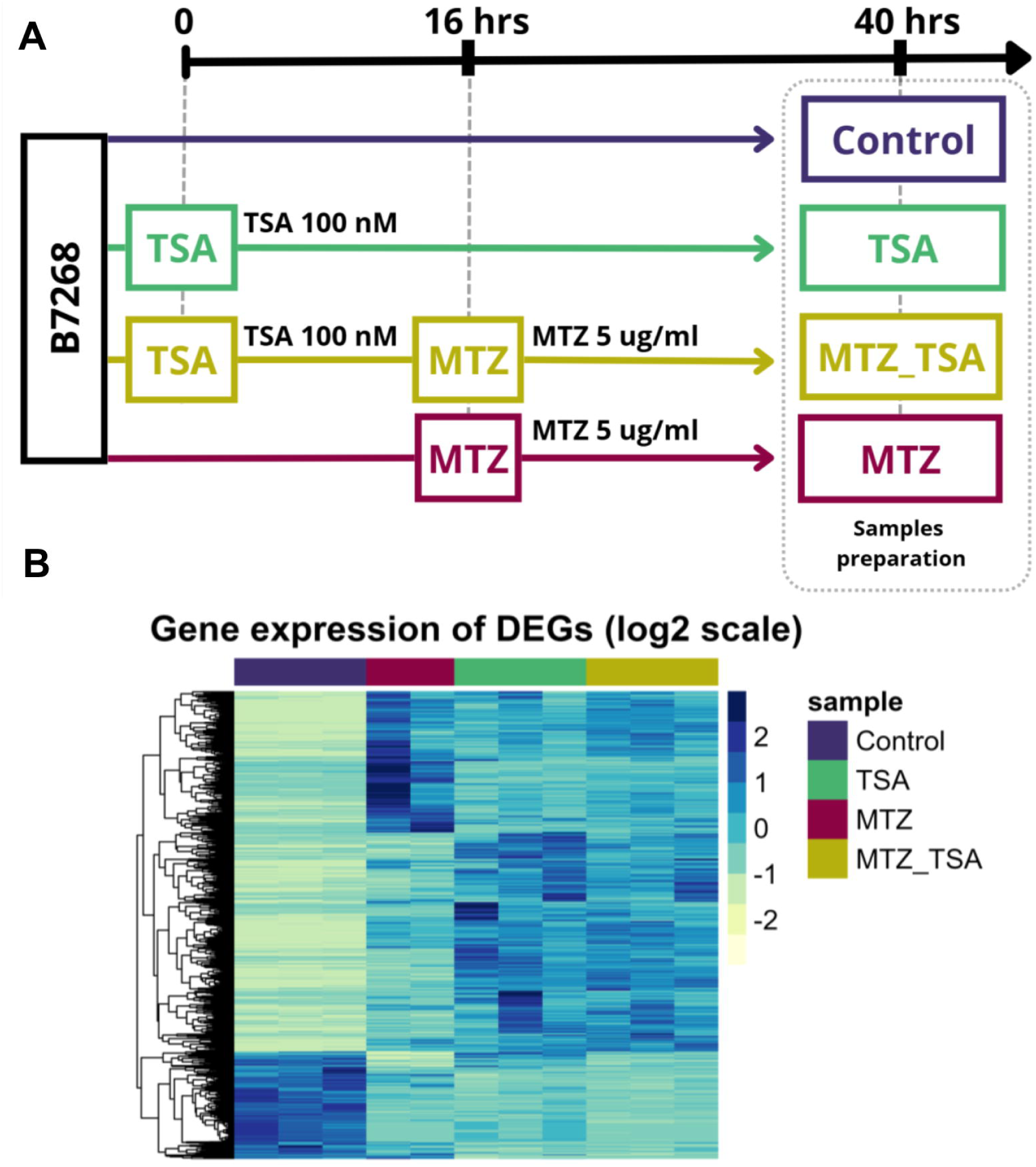
A. RNA-seq experimental Design. The RNA-seq experiment was designed with four treatment groups: (1) untreated B7268 parasites control, (2) B7268 was treated with 100 nM Trichostatin A (TSA), (3) B7268 was treated with 5 μg/ml metronidazole (MTZ) and (4) B7268 was treated with 100 nM TSA for 16 h followed by a treatment with 5 μg/ml metronidazole (TSA_MTZ). All samples, including controls, were prepared for sequencing at the same time, 40 hours after the start of the experiment. B. Heatmap displaying normalized expression of DEGs obtained throughout the different treatments. Each horizontal line represents an individual gene. Color gradient represents the expression level.

As increased histone acetylation results in a more relaxed chromatin state that promotes gene expression [30], we focused on identifying genes potentially associated with MTZ resistance whose expression was upregulated by TSA-induced changes. We selected two groups of upregulated genes: (1) genes upregulated in both TSA and TSA_MTZ treatments (1172 genes) and (2) genes exclusively upregulated in TSA_MTZ treatment (292 genes), resulting in 1464 TSA-associated induced genes (Fig. 3D and Supplementary Table 2). Gene Ontology (GO) enrichment analysis of these genes revealed significant overrepresentation of terms related to cell adhesion, DNA-binding transcription factor activity, and peptidase activity, as well as genes involved in transmembrane transport processes, such as drug transmembrane transporters (Fig. 3E). Functional categorization of these genes (Supplementary Table 2) showed that the largest group comprised genes with an unspecified product (301 genes). Among genes with predicted functional domains, prominent categories included those associated with ubiquitin-related processes and the proteasome system (204 genes), kinases and phosphatases (117 genes), transcriptional regulators (78 genes), pathogenesis-related proteins (58 genes), transporters (25 genes), redox-related proteins (18 genes), and chromatin-associated factors (8 genes), among others (Supplementary Table 2). Previously identified genes involved in MTZ activation, such as Thioredoxin Reductase (TVAGG3_0315040) and Nitroreductases (TVAGG3_0444150 and TVAGG3_0873820), were significantly upregulated in TSA_MTZ treated parasites, supporting their role in MTZ sensitivity (Supplementary Table 2). Furthermore, genes from the Diflavin Flavoprotein A 2-related family, Flavodoxin-like fold, and Flavoprotein family (TVAGG3_0421620, TVAGG3_0583440, TVAGG3_0080430) were also upregulated in MTZ_TSA group. Notably, redox-related proteins like Flavodoxins and hydrogenosomal enzymes showed varied expression, suggesting a complex interplay in resistance mechanisms. As example, a Ferrodoxin (Fd) and three PFOR genes (TVAGG3_0476390, TVAGG3_0282970, TVAGG3_0890230, TVAGG3_0998520) were downregulated in both MTZ_TSA and TSA groups, suggesting that transcription of these genes does not seem to be regulated by histone acetylation (Supplementary Table 2). To validate the RNA-seq results, quantitative real-time PCR (qPCR) was performed on selected genes exhibiting TSA responsive expression changes. Four genes induced by TSA_MTZ treatment were selected: TVAGG3_0172050, TVAGG3_1037610, TVAGG3_0371210, and TVAGG3_0315040 (thioredoxin reductase). In agreement with the RNA-seq data, qPCR analysis confirmed significant upregulation of these genes in TSA-treated B7268 parasites compared to untreated controls (Fig. 3F). Collectively, these results validate the transcriptomic impact of TSA and support the role of histone acetylation in modulating the expression of genes potentially involved in MTZ resistance mechanisms.

**Figure 3:**
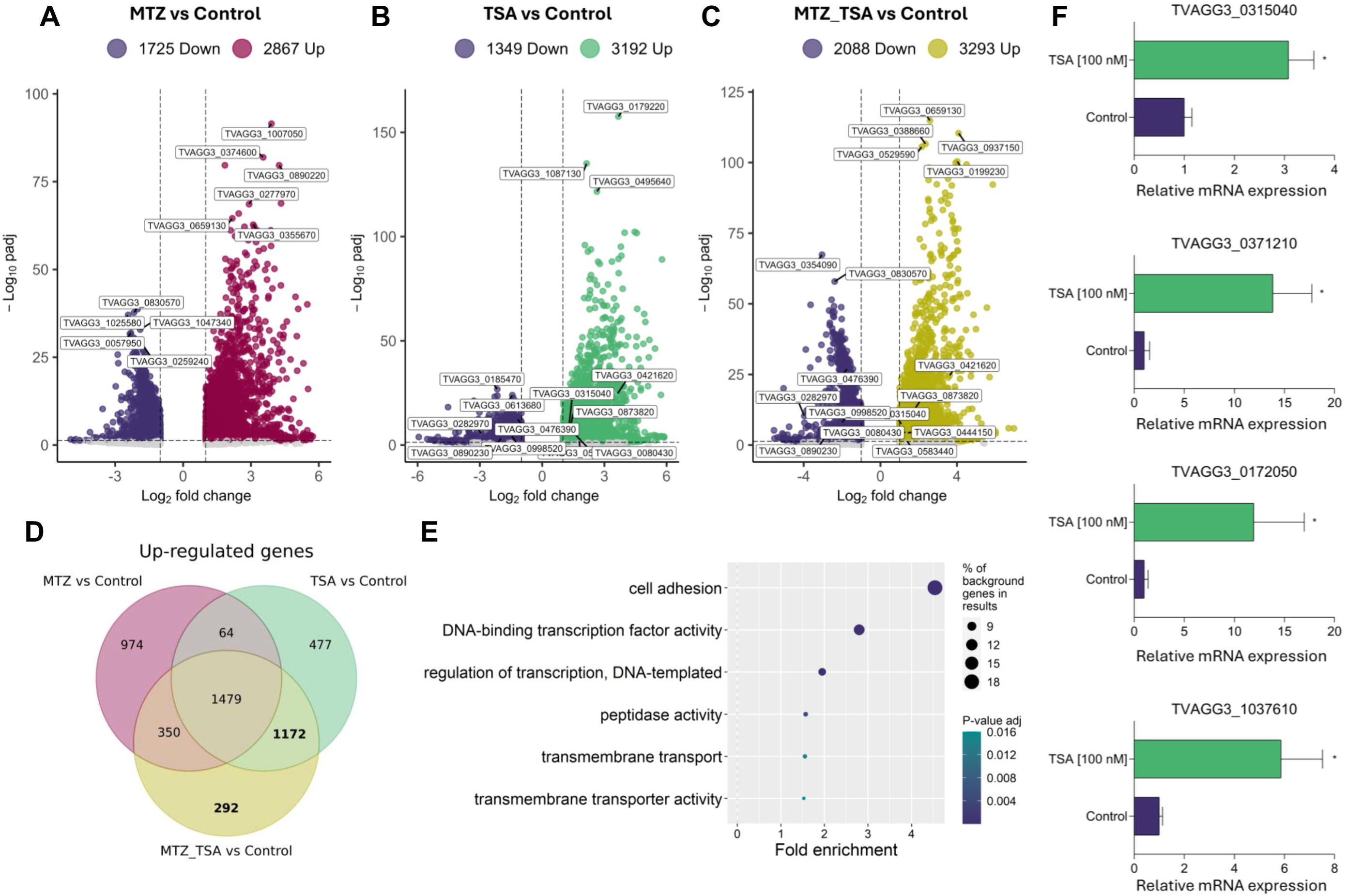
Volcano plot distributions of differentially expressed genes (DEGs) obtained from RNA-seq analysis of B7268 treated with MTZ vs Control (A), B7268 treated with TSA vs Control (B) and B7268 treated with MTZ + TSA vs Control (C). The selective condition to determine DEGs was defined as |log2FC| > 1 and p-value adjusted < 0.05. (D) Venn diagram showing the number of up-regulated genes in each treatment as well as the shared by the three comparisons. The genes of interest are highlighted in bold. (E) Gene ontology enrichment analysis of molecular function for genes of interest. (F) qPCR validation of TSA-induced gene expression. Relative expression levels of four genes upregulated by TSA treatment in B7268 parasites: TVAGG3_0172050 (protein of unknown function), TVAGG3_1037610 (nucleotide-diphospho-sugar transferases family), TVAGG3_0371210 (glycosyl hydrolase) and TVAGG3_0315040 (thioredoxin reductase). Parasites were treated with TSA (100 nM) or left untreated (Control). Expression values are normalized to the housekeeping gene Hmp24. Data is expressed as fold increase compared with the control sample ± SD. * = P-value ≤ 0.05 (Student’s t-test).

### Genome-wide mapping of H3K9Ac displays functional role in gene activation and modulation by TSA treatment

To investigate the direct contribution of histone acetylation to chromatin remodeling associated with TSA and MTZ resistance modulation, we performed ChIP-seq to map the genome-wide distribution of the activating histone mark H3K9Ac in the highly resistant B7268 strain under the same conditions used for RNA-seq analysis: untreated control and sequential TSA followed by MTZ treatment (MTZ_TSA). Principal component analysis (PCA) revealed that replicate samples cluster more closely with each other than with samples from different treatments, with PC1 and PC2 explaining 58% and 31% of the variance, respectively. (Supplementary Fig. 2). Genome-wide distribution of H3K9Ac observed in our study exhibits a bimodal pattern across promoter regions and gene bodies, with a predominant enrichment within gene bodies rather than at promoters (Fig. 4A and Supplementary Fig. 3). This recapitulates the pattern reported for H3K27Ac in *T. vaginalis* by Song et al. ^28^; suggesting that active histone acetylation marks in this parasite may follow a gene body enriched deposition pattern, distinct from the promoter-proximal enrichment typically observed in other eukaryotic models. Comparative analysis of H3K9Ac profiles across treatment conditions showed that untreated parasites characterized by narrow, sharply defined peaks centered around transcription start sites (TSS) (Fig. 4A and Supplementary Fig. 3). In contrast, parasites subjected to sequential MTZ_TSA treatment exhibited broader H3K9Ac peaks, consistent with increased histone acetylation resulting from HDAC inhibition (Fig. 4A). Following peak calling, 10,135 peaks were identified in control samples and 5,394 in MTZ_TSA-treated parasites (Supplementary Table 3).

**Figure 4:**
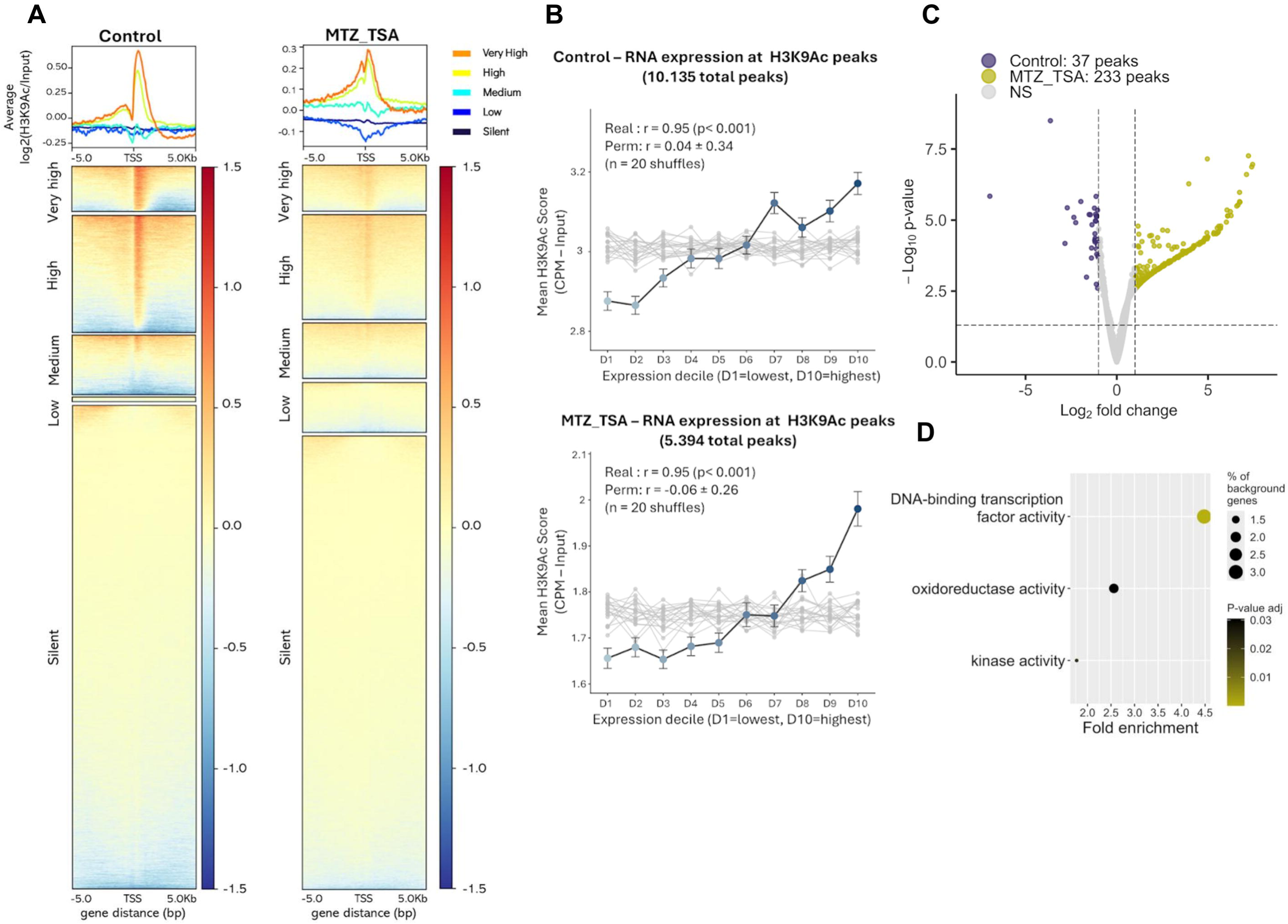
A. H3K9Ac ChIP-seq signal profiles around the transcription start sites (TSS) in B7268 parasites under Control and MTZ_TSA (100 nM TSA + 5 µg/ml MTZ) conditions. Genes are stratified by RNA-seq expression levels (Very high: RPKM ≥1000; High: 100≤RPKM<1000; Medium: 10≤RPKM<100; Low: 1≤RPKM<10; Very low: RPKM<1). Heatmaps show log_2_ fold-enrichment over input normalized signal ±5 kb from TSS. Average profiles are displayed above each heatmap. B. Mean H3K9Ac score (ChIP CPM – Input CPM) per expression decile for genes intersecting with H3K9Ac peaks in each condition. Genes were ranked by mean RNA-seq expression, partitioned into ten equal-sized deciles (D1 to D10) and ChIP signal score was averaged across replicates. Each point represents the mean ± SEM of H3K9Ac scores within a decile. The monotonic association between expression rank and ChIP score was quantified using the Spearman rank correlation coefficient. Gray lines represent 20 random permutations of expression labels; the mean ± SD of permuted Spearman *r* values is reported alongside the real correlation coefficient as a negative control. C. Volcano plots of differential H3K9Ac peak analysis in MTZ_TSA vs control samples. Colored points indicate significantly enriched peaks (padj < 0.05, |log □FC| > 1). Peak numbers shown in each category. D. Gene Ontology enrichment analysis of genes associated with MTZ_TSA-enriched peaks (n=233). Dot size represents percentage of background genes, and color intensity indicates adjusted p-value. Only significantly enriched molecular function terms are shown (padj < 0.05).

Integration of ChIP-seq and RNA-seq data revealed that H3K9Ac enrichment is strongly associated with transcriptional activity. Genes with “very high” expression levels showed the highest H3K9Ac signal intensity across all treatment conditions, while genes with “low” or “silent” expression exhibited minimal or absent H3K9Ac enrichment across all treatment conditions (Fig. 4A). To further quantify this relationship, genes intersecting with H3K9Ac peaks were ranked by their DESeq2 normalized mean expression and partitioned into ten equal sized deciles (D1 = lowest expression, D10 = highest expression), and the mean ChIP score per decile was computed and correlated with expression rank using the Spearman rank correlation. In both conditions, mean H3K9Ac score increased monotonically from the lowest to the highest expression decile, with correlation coefficients of r = 0.95 for control and MTZ_TSA treated parasites (p < 0.001 in both cases; Color line in Fig. 4B). To confirm that these correlations reflect a genuine biological relationship rather than a statistical artifact of the decile grouping procedure, expression labels were randomly permuted 20 times and the full analysis was repeated for each shuffle. The resulting permuted correlations clustered around zero (r = 0.04 ± 0.34 and −0.06 ± 0.26 for Control and MTZ_TSA samples, respectively. Gray lines in Fig. 4B). When this analysis was extended to all annotated protein coding genes genome-wide, expressed genes displayed a progressive increase in ChIP signal with expression level in all cases (Supplementary Fig. 4). Together, these results demonstrate a robust and genome-wide positive association between H3K9Ac levels and gene expression, supporting a biologically meaningful link between this histone mark and transcriptional activity in T. vaginalis.

To identify H3K9Ac enriched regions specifically associated with TSA-mediated modulation of MTZ resistance, we performed differential peak analysis comparing MTZ_TSA treatment to untreated control. This analysis revealed 233 peaks enriched in MTZ_TSA-treated parasites relative to control (Fig. 4C and Supplementary Table 4). Gene Ontology enrichment analysis of genes associated with the 233 differential peaks in MTZ_TSA treated parasites identified significant enrichment of terms related to DNA-binding transcription factor activity, oxidoreductase activity and kinase activity (Fig. 4D). These findings suggest that TSA-induced histone acetylation might be reshaping the chromatin landscape to modulate transcriptional regulation and redox homeostasis, processes likely contributing to altered MTZ susceptibility ^53^.

### Cross-strain transcriptomic validation identifies conserved epigenetically regulated MTZ sensitivity genes

Given our hypothesis that epigenetic silencing contributes to MTZ resistance, we focused on genes downregulated in resistant strains relative to sensitive ones. To identify candidate genes that may be epigenetically silenced in resistance but reactivated by histone acetylation, we intersected the 1,464 genes induced by TSA and MTZ treatment (Fig. 3D) with the 1,857 genes upregulated in the MTZ-sensitive strain NYH209 in our RNA-seq dataset ^33^ (Supplementary Table 5), as well as genes consistently upregulated across nine publicly available RNA-seq datasets from additional MTZ sensitive strains (G3, GOR69, NYCA04, NYCB20, NYCD15, NYCE32, NYCF20, NYCG31, and SD2), all in comparison to the MTZ-resistant strain B7268 (accessions SRP057357 and SRP057311) ^15^ (Fig. 5A). Genes were considered differentially expressed in the multi-strain analysis if they were significantly expressed in at least three of the nine MTZ sensitive strains and not upregulated in the resistant strain B7268 in any comparison with these sensitive strains. This integrative analysis identified 66 high confidence candidate genes that are expressed in sensitive strains and become induced upon TSA treatment (Fig. 5A and Supplementary Table 6), suggesting that their repression in B7268 is epigenetically mediated trough histone acetylation. Analysis of these 66 genes revealed enrichment in functional categories related to surface antigens, cytoskeleton, transcription regulation, metabolism redox stress processes, among others (Fig. 5B). Representative genes from this group, including thioredoxin reductase (TVAGG3_0315040), an oxidoreductase (TVAGG3_0792210), a cyclin-dependent kinase inhibitor (TVAGG3_0529700) and the transcription factor Myb6 (TVAGG3_0427690), showed low basal expression in untreated B7268 resistant parasites and in parasites exposed to MTZ alone (Fig. 5C). In contrast, treatment with TSA (100 nM) or sequential TSA followed by MTZ resulted in robust transcriptional upregulation of these genes, as evidenced by increased RNA-seq coverage (Fig. 5C), indicating that their activation is linked to histone acetylation-mediated chromatin remodeling. Consistently, the expression of these genes was higher in the naturally sensitive NYH209 strain, as well as in other previously reported MTZ-sensitive strains, compared with the highly resistant B7268 strain (Fig. 5C). Together, these findings suggest that the expression of these genes is epigenetically regulated in resistant parasites and may contribute to MTZ resistance.

**Figure 5:**
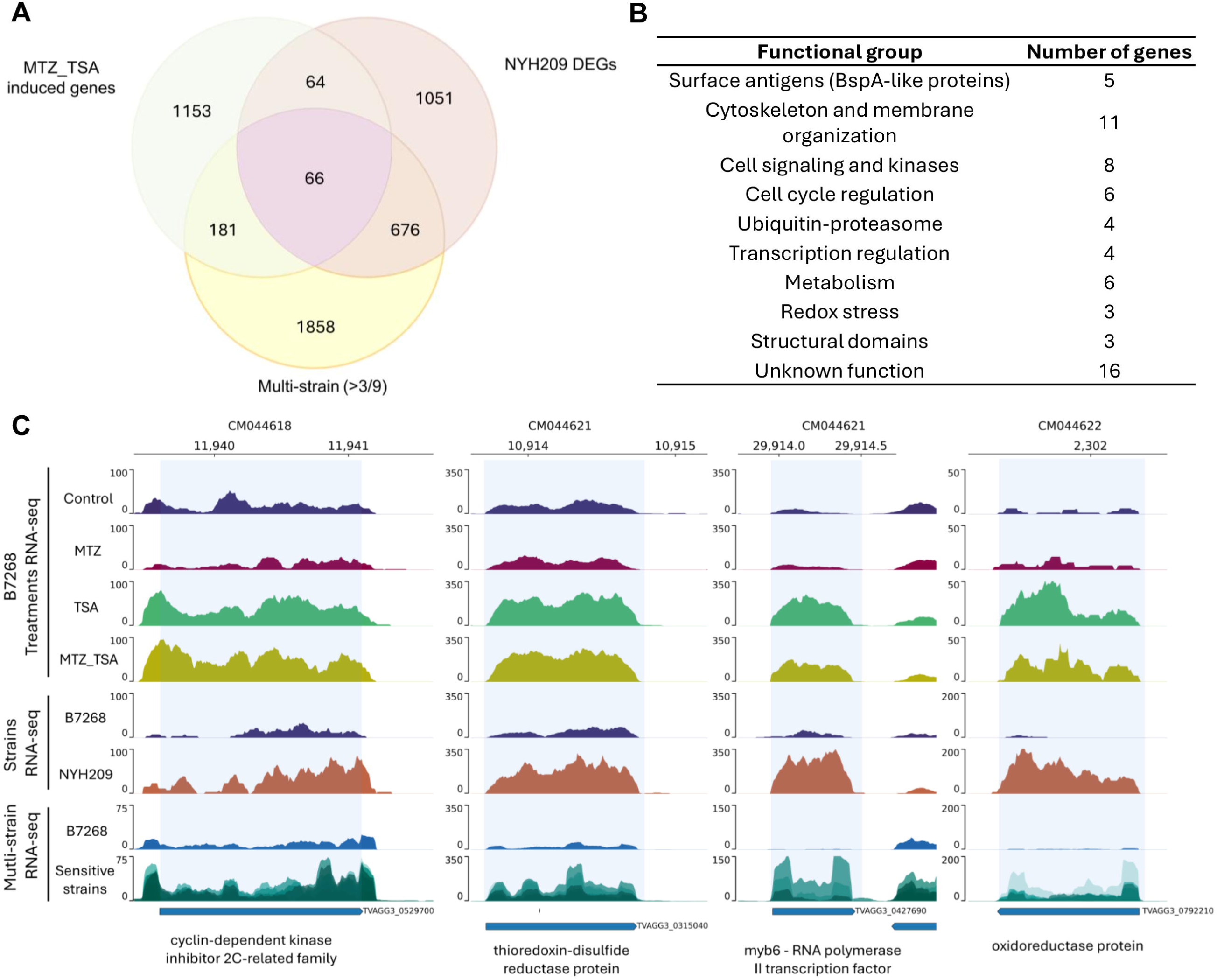
A. Venn diagram illustrating the overlap among three gene sets: MTZ_TSA-induced genes in B7268 (1464 genes), differentially expressed genes in sensitive strain NYH209, and genes consistently differentially expressed across multiple sensitive strains (DEGs in at least 3 of 9 strains analyzed). The intersection of all three datasets yielded 66 high-confidence candidate genes epigenetically modulated genes that might be involved in MTZ resistance mechanisms. B. Table summarizing the functional groups of the 66 high-confidence candidate genes. C. Integrated genomic tracks showing RNA-seq profiles of four representative high-confidence candidate genes: thioredoxin reductase (TVAGG3_0315040), oxidoreductase protein (TVAGG3_0792210), cyclin-dependent kinase inhibitor 2C-related family (TVAGG3_0529700), and transcription factor Myb6 (TVAGG3_0427690). Top tracks: RNA-seq coverage in B7268 parasites under Control, MTZ (5 µg/ml), TSA (100 nM), and MTZ_TSA treatments. Middle tracks: RNA-seq coverage comparing B7268 (resistant) and NYH209 (sensitive) strains. Bottom tracks: Multi strains RNA-seq coverage comparing B7268 strain against 9 different sensitive strains (G3, GOR69, NYCA04, NYCB20, NYCD15, NYCE32, NYCF20, NYCG31, SD2). Y-axis shows normalized read counts; genomic coordinates are indicated above each gene. Gray shading highlights gene body regions.

### Epigenetically regulated thioredoxin-disulfide reductase and a Myb-like containing domain gene modulate MTZ resistance in a highly resistant isolate

To functionally validate the contribution of epigenetically silenced genes to MTZ resistance in the B7268 strain, we selected two candidates out of the 66 genes set for targeted overexpression: Thioredoxin-disulfide reductase (TrxR, TVAGG3_0315040), and a Myb-like containing domain gene (Myb6, TVAGG3_0427690). These candidates were prioritized based on (i) the relevance of redox homeostasis and transcriptional regulation to drug resistance mechanisms, and (ii) their consistent upregulation across multiple MTZ sensitive strains relative to B7268 (Fig. 6A). Notably, both genes exhibited significant overexpression (log2FC > 1) in at least three independent sensitive strains (Fig 6A). To assess whether restoring expression of these genes could reverse the resistant phenotype, each gene was independently cloned into episomal expression vectors under the control of a strong promoter and expressed as C-terminal HA-tagged fusion proteins in the highly resistant B7268 background. Immunofluorescence assays showed that Myb6-HA was localized predominantly to the nucleus and TrxR-HA exhibited perinuclear distribution (Fig. 6B). Functional assessment of MTZ susceptibility in the transfectants showed that Myb6 overexpression significantly reduced resistance, decreasing the minimum inhibitory concentration (MIC) from 168 µg/ml to 32 µg/mL (p = 0.0078) compared to parasites transfected with empty vector control (EpNeo). Similarly, TrxR overexpression resulted in a significant reduction in resistance, with the MIC decreasing to 51.8 µg/ml (p = 0.0431) (Fig. 6C). Together, these results provide direct functional evidence that TrxR and Myb6 contribute to MTZ sensitivity and suggest that their reduced expression in the B7268 strain might be a key determinant of the resistant phenotype. These results support a model in which epigenetic silencing of specific genes contributes to MTZ resistance and can be reversed through their reactivation.

**Figure 6:**
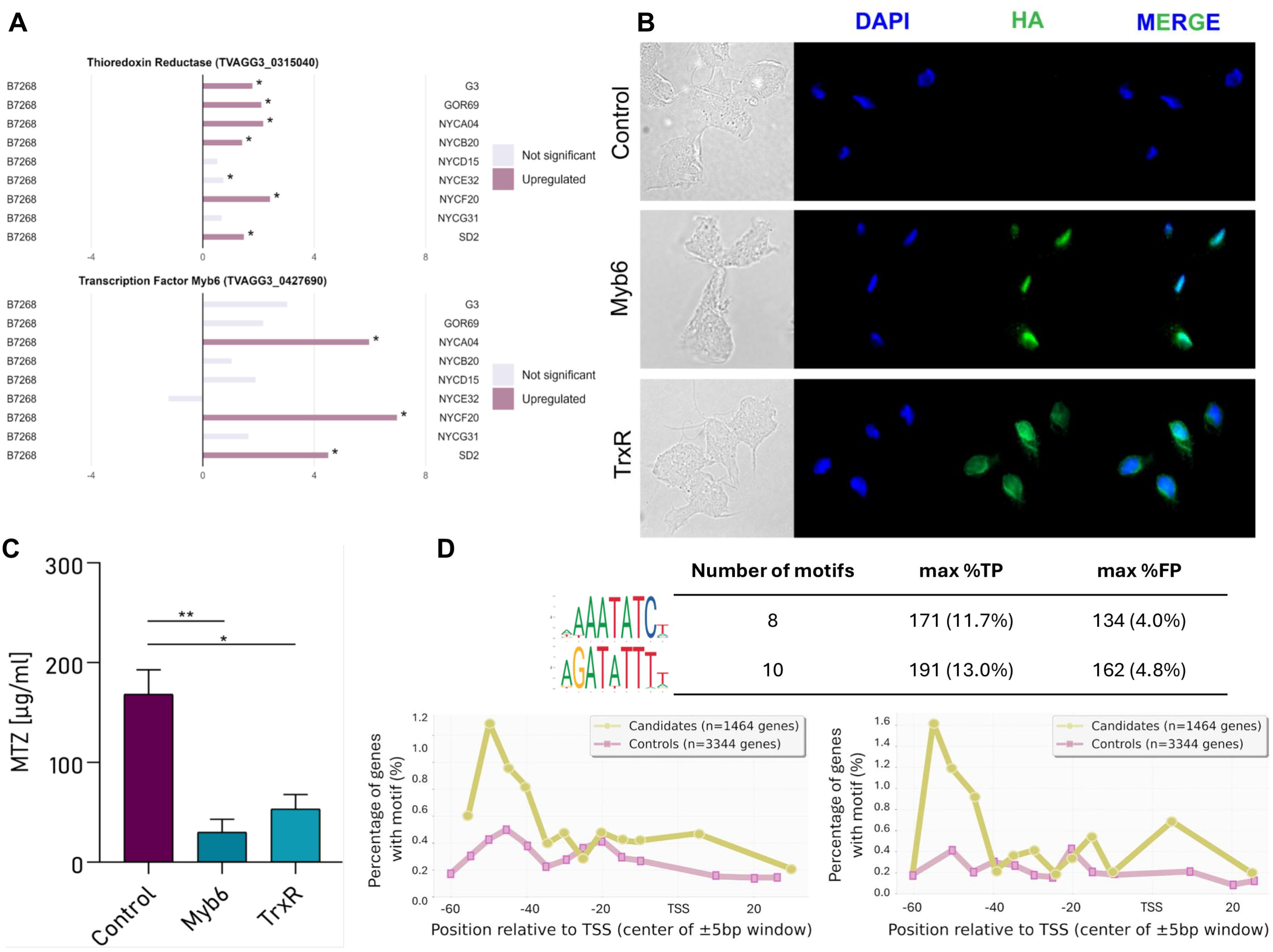
A. Expression profiles of thioredoxin reductase (TVAGG3_0315040) and transcription factor Myb6 (TVAGG3_0427690) candidate genes across nine MTZ-sensitive strains (G3, GOR69, NYCA04, NYCB20, NYCD15, NYCE32, NYCF20, NYCG31, SD2) and the resistant B7268 strain. Log_2_ fold-change values show gene expression in each sensitive strain relative to B7268. Bars are colored by differential expression significance: upregulated (pink) or not significant (gray). Asterisks indicate statistically significant differences (* = padj < 0.05). Data derived from reanalysis of public datasets (SRP057357 and SRP057311). B. Subcellular localization of two selected putative genes involved in modulation of drug resistance: thioredoxin-disulfide reductase (TrxR, TVAGG3_0315040) and Myb-like containing domain (Myb6, TVAGG3_0427690) transfected in B7268 strain. Cells expressing C-terminal HA-tagged versions of the indicated gene were stained for immunofluorescence microscopy using a mouse HA-tagged antibody. The nucleus (blue) was also stained with 4-,6-diamidino-2-phenylindole. C. The level of MTZ resistance of parasites from the B7268 strain transfected with TrxR and Myb6 was assessed and compared with parasites transfected with control empty plasmid. Four independent experiments were performed. Data are expressed as mean values ± SD of three independent experiments. D. Enrichment analysis of AATATC and AGATAT-core Myb binding motifs. Top: consensus sequence logos from significantly enriched Myb-related PWMs (MA1187.2 and MA1184.1 shown as representative). The table shows motif statistics: number of PWMs with each pattern, maximum true positive rate (candidate genes, n=1,464) and false positive rate (control genes, n=3,344). Bottom: positional conservation plots (MA1187.2 and MA1184.1) displaying motif enrichment across promoter regions (−60 to +40 bp relative to TSS, center of ±5 bp window). Both motif patterns show significant enrichment peaks ∼60 bp upstream of TSS in candidate gene promoters.

### Enrichment of Myb binding motifs in promoters of TSA-induced genes supports a role for Myb6 in modulating gene expression associated with drug resistance

Given that the putative transcription factor Myb6 was functionally validated as a modulator of MTZ sensitivity in the resistant B7268 strain, we next examined whether Myb binding motifs were enriched within promoter regions of TSA_MTZ induced genes. To this end, we retrieved 261 Myb-related position weight matrices (PWMs) from the JASPAR database and performed motif enrichment analysis using AME (Analysis of Motif Enrichment). To this end, promoter sequences (−60 to +40 bp relative to TSS) from the 1,464 MTZ_TSA-induced genes were compared with a background set of 3,344 genes that were also upregulated in these experiments but were not part of the MTZ_TSA induced gene set (Fig. 3D). AME analysis revealed significant enrichment of Myb-related motifs in the promoters of MTZ_TSA induced genes following Bonferroni correction for multiple testing. Despite originating from distinct JASPAR entries, the highest-ranking motifs clustered into two conserved sequence patterns (Fig. 6D). The first group centered on an AATATC core motif and included 8 closely related PWMs (MA2353.1, MA0972.1, MA1184.2, MA1184.1, MA1191.1, MA1191.2, MA1401.2, and MA1401.1) (Fig. 6D). The second group was defined by an AGATAT core motif and comprised 10 highly similar PWMs (MA1187.2, MA1183.2, MA1187.1, MA1185.2, MA1183.1, MA1182.2, MA1190.2, MA1182.1, MA1185.1, and MA1190.1) (Fig. 6D).

Positional conservation analysis revealed highly consistent localization of both motif groups, with a prominent enrichment peak approximately 50 bp upstream of the TSS (Fig. 6D). Representative motifs are shown for clarity (MA1187.2 for the AGATAT pattern and MA1184.1 for the AATATC pattern) (Fig. 6D). Quantitative analysis demonstrated that AGATAT motifs were present in 13.0% of MTZ_TSA induced genes, compared with 4.8% of the genes in the control set (Fig. 6D and Supplementary Table 7). Similarly, AATATC-pattern motifs were detected in 11.7% induced genes versus 4.0% control genes (Fig. 6D and Supplementary Table 7). Together, these findings suggest that TSA-induced chromatin remodeling promotes the binding of Myb6 at defined regulatory positions flanking the TSS likely coordinating expression of genes involved in MTZ resistance modulation.

## DISCUSSION

Metronidazole (MTZ) resistance in Trichomonas vaginalis is a multifactorial process involving partially overlapping pathways that affect drug activation, redox balance, and energy metabolism. Clinical metronidazole resistance in *T. vaginalis* has been linked to reduced flavin reductase activity, an enzyme that removes intracellular oxygen ^12^. Loss of this activity leads to elevated intracellular oxygen levels, promoting futile cycling and detoxification of activated MTZ, thereby reducing drug efficacy ^12^. In addition, decreased activity of enzymes directly involved in drug activation, thioredoxin reductase (TrxR) and flavin reductase, further contributes to reduced drug efficacy ^9,12^. While this oxygen dependent (“aerobic”) resistance mechanism is frequently observed in clinical isolates ^10,11^, laboratory-induced resistance is often linked to impaired hydrogenosomal function, including downregulation of ferredoxin and pyruvate:ferredoxin oxidoreductase ^10^. Consistent with these biochemical alterations, transcriptomic analyses revealed widespread gene expression changes in MTZ resistant *T. vaginalis* isolates ^15,33^, and similar associations between MTZ resistance and transcriptional reprogramming have been reported in the related parasite *Entamoeba histolytica* ^54^. Complementing these findings, genome-wide association studies have identified 72 single nucleotide polymorphisms associated with MTZ resistance, including variants in pyruvate:ferredoxin oxidoreductase ^15^ and nitroreductase genes ^14^. Our findings add an additional regulatory dimension by demonstrating that epigenetic mechanisms directly contribute to MTZ resistance. Inhibition of histone deacetylases (HDACs) with trichostatin A (TSA) induced widespread transcriptional changes, increased H3K9 acetylation, and enhanced drug sensitivity in a highly resistant strain. The positive association between H3K9 acetylation and transcriptional activation supports a model in which chromatin remodeling facilitates a more permissive transcriptional state. These results extend previous observations linking histone acetylation and DNA methylation to gene regulation in *T. vaginalis* ^26,28,33,55,56^, and align with studies in other systems where chromatin modifiers influence antimicrobial responses. Similar effects have been observed with Resveratrol, a natural polyphenolic compound known to elicit diverse epigenetic modifications such as alterations in DNA methyltransferases, HDACs, and lysine-specific demethylase ^57^. This compound has previously demonstrated its ability to induce antiparasitic activity through the modulation of gene expression and hydrogenosomal dysfunction in *T. vaginalis* ^58^. Likewise, chromatin modifiers have directly implicated in the ability of *Candida spp.* to survive in the presence of anti-fungal drugs ^59^. For instance, the deletion of the Rpd3/Hda1 family of histone deacetylases in *C. albicans* sensitizes the yeast to fluconazole, itraconazole, and voriconazole ^60^. Consistently, the removal of the Histone Acetyltransferase Hat1 achieves the reverse, increasing *C. albican*’s azole tolerance ^61^. Together, these parallels highlight a conserved role for chromatin-based regulatory mechanisms in shaping antimicrobial responses.

Building on this framework, we propose that epigenetic silencing contributes to the resistant phenotype. TSA treatment of the resistant B7268 strain led to the upregulation of 1,464 genes, including those involved in transcriptional regulation and redox processes. Integration of these TSA-responsive genes with expression profiles from MTZ-sensitive strains identified 66 high-confidence candidates that are consistently expressed in sensitive backgrounds but repressed in B7268. Functional validation of two candidates, TrxR and the Myb-like transcription factor Myb6, demonstrated that their overexpression partially restores MTZ susceptibility, indicating that reduced expression of these genes contributes directly to resistance. These results highlight the importance of combining integrative transcriptomic analyses with functional validation to identify genes that play a causal role in MTZ resistance.

The role of redox metabolism in MTZ resistance is well established. A strong link between MTZ resistance and redox metabolism has been established in both *Entamoeba histolytica* and *T. vaginalis* ^53^. In *T. vaginalis*, the thioredoxin-peroxiredoxin system constitutes a central antioxidant pathway ^62–64^, with TrxR maintaining redox homeostasis and supporting peroxiredoxin function ^53^. In this context, our data provide functional evidence that TrxR contributes to MTZ resistance, as its overexpression in the resistant strain significantly reduced drug resistance. Notably, TrxR has previously been proposed as a direct target of 5-nitroimidazoles, which can form covalent adducts with cysteine residues in the enzyme, thereby inhibiting its activity ^9^. Indeed, total insensitivity to metronidazole can be rapidly induced in *T. vaginalis* by treatment with diphenylene iodonium (DPI), which inhibits all flavoenzymes in the parasite including TrxR, thereby blocking the activation of MTZ ^65^. Similarly, nitroreductases (ntr4) have presented mutations associated with the development of resistance in *T. vaginalis* parasites ^66^. In agreement with these observations, our RNA-seq analysis identified two nitroreductase genes whose expression was modulated by TSA and MTZ treatment, further linking epigenetic regulation to known resistance-associated pathways.

Beyond redox enzymes, our data identify Myb6 as a previously unrecognized regulator of MTZ resistance in *T. vaginalis*. Although Myb genes have been previously described as regulators of gene expression in this parasite ^67–70^, they have not been associated with drug resistance. Motif enrichment analysis revealed a significant overrepresentation of Myb-binding sites in promoters of TSA-induced genes, with a conserved positional bias consistent with direct transcriptional regulation. Together with reports of MYB dysregulation in resistant parasites ^71^, and their roles in drug response in other systems ^72^, these findings suggest that Myb6 may coordinate a broader transcriptional program governing MTZ susceptibility. Future studies should aim to define the Myb6 regulon and determine whether simultaneous reactivation of multiple epigenetically silenced genes further enhances MTZ sensitivity.

Collectively, our results support a model in which epigenetic regulation, particularly histone acetylation, modulates the expression of genes involved in MTZ activation and resistance. This highlights HDACs as promising therapeutic targets for the treatment of MTZ-resistant trichomoniasis. Similar mechanisms have been described in other protozoan parasites, such as *Plasmodium falciparum* and *Trypanosoma spp*., where chromatin-based regulation influences drug response and persistence ^73–75^. In *P. falciparum*, HDAC-mediated silencing of drug transport genes reduces drug uptake ^76^ while in *T. brucei*, they regulate telomeric silencing of variant surface glycoprotein (VSG) expression sites, facilitating immune evasion and resistance ^77^. Notably, blasticidin S-resistant *P. falciparum* parasites silence the clag2 and clag3 genes (cytoadherence linked antigen genes) through loss of the activating histone marks H3K9Ac and H3K4me3 modifications, reducing channel-mediated antibiotic uptake ^78^. Importantly, the reversible nature of epigenetic modifications highlights their therapeutic potential. Consistently, HDAC inhibitors, including TSA and suberanilohydroxamic acid (SAHA), have been shown to induce global histone hyperacetylation, reactivating silenced genes and disrupting resistance mechanisms in *P. falciparum* ^73,74,79^. However, evidence from other protozoan systems suggests that durable modulation of resistance-related gene expression likely requires coordinated interactions between multiple epigenetic pathways, including histone acetylation, methylation, and chromatin remodeling ^73,74,79^. A deeper understanding of these mechanisms will be essential for the development of more effective strategies to combat MTZ resistant infections

## Acknowledgments

We thank our colleagues in the laboratory for helpful discussions and Maria Florencia Irigoyen and Carlos Alberici for their technical assistance. This research was supported with a grant from the ANPCyT grant BID PICT-2018-01892 (NdM), an NIH grant (5R01AI160387) (NdM), an International Union of Biochemistry and Molecular Biology (IUBMB) Wood-Whelan Research Fellowship (to D.M.) and a Company of Biologists Travelling Fellowship (to J.S.). Additional support was provided by the National Institute of Allergy and Infectious Diseases (NIAID) [R01 AI136511, R01 AI188634, and R21 AI142506 to K.L.R.], and by the University of California, Riverside (NIFA-Hatch-225935 to K.L.R.). .P.H.S-M., V.C. and N.dM. are researchers from the National Council of Research (CONICET) and National University of San Martin (UNSAM). D.M. and L.I are PhD fellow from CONICET and J.S is a PhD fellow from ANPCyT. The funders had no role in study design, data collection and analysis, decision to publish, or preparation of the manuscript.

## DATA AVAILABILITY

All data supporting the findings of this study are available within the paper and its supplementary Information. All sequencing data that support the findings of this study have been deposited in the National Center for Biotechnology Information Sequence Read Archive (SRA) and are accessible through accession number PRJNA1215954.

