## Supplementary figures and images for "Epigenetic reprogramming restores metronidazole sensitivity in drug-resistant *Trichomonas vaginalis* parasites"

### Supplementary Figures

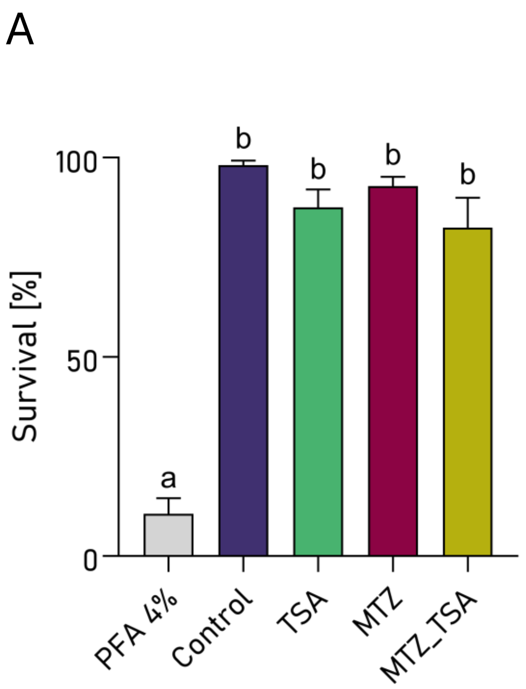

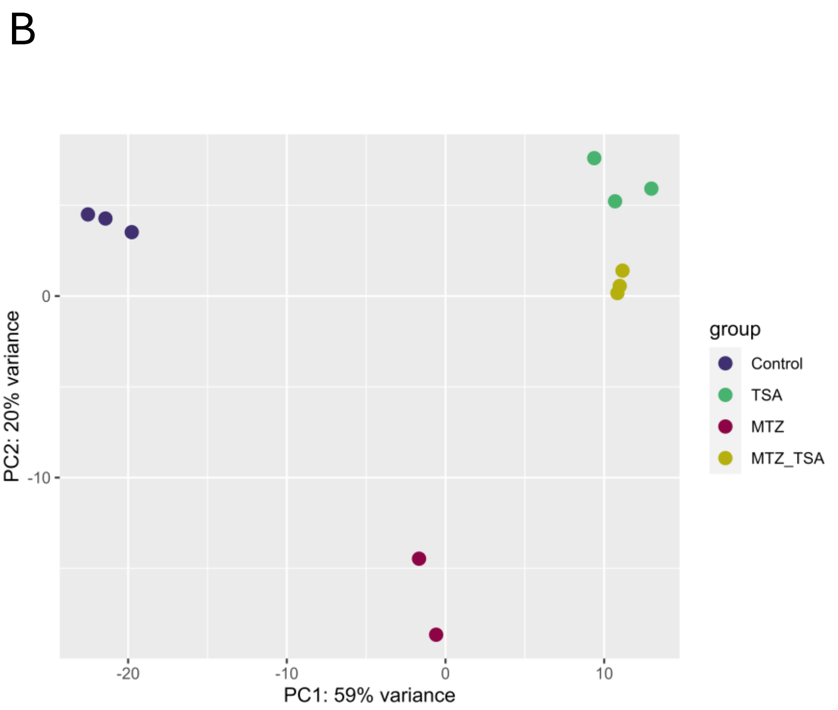

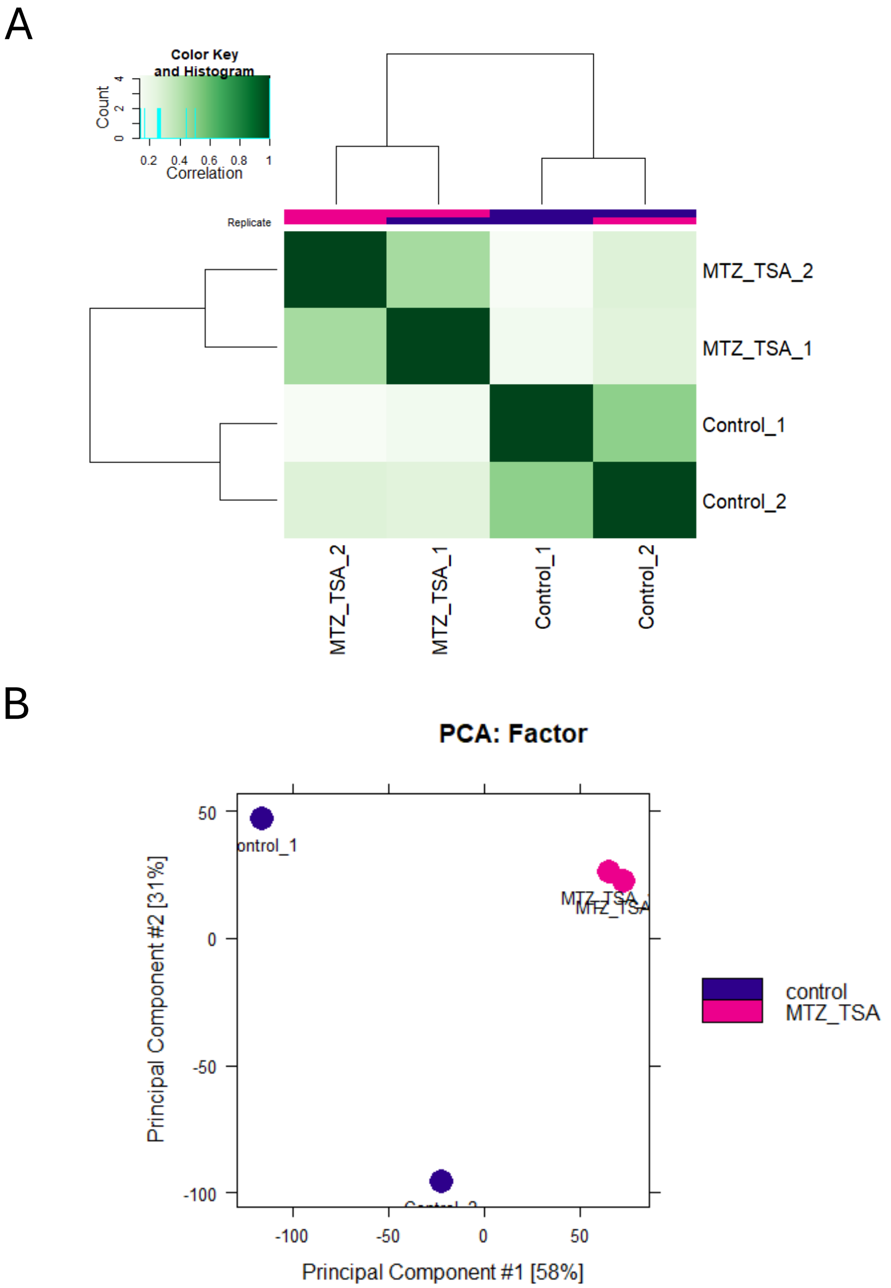

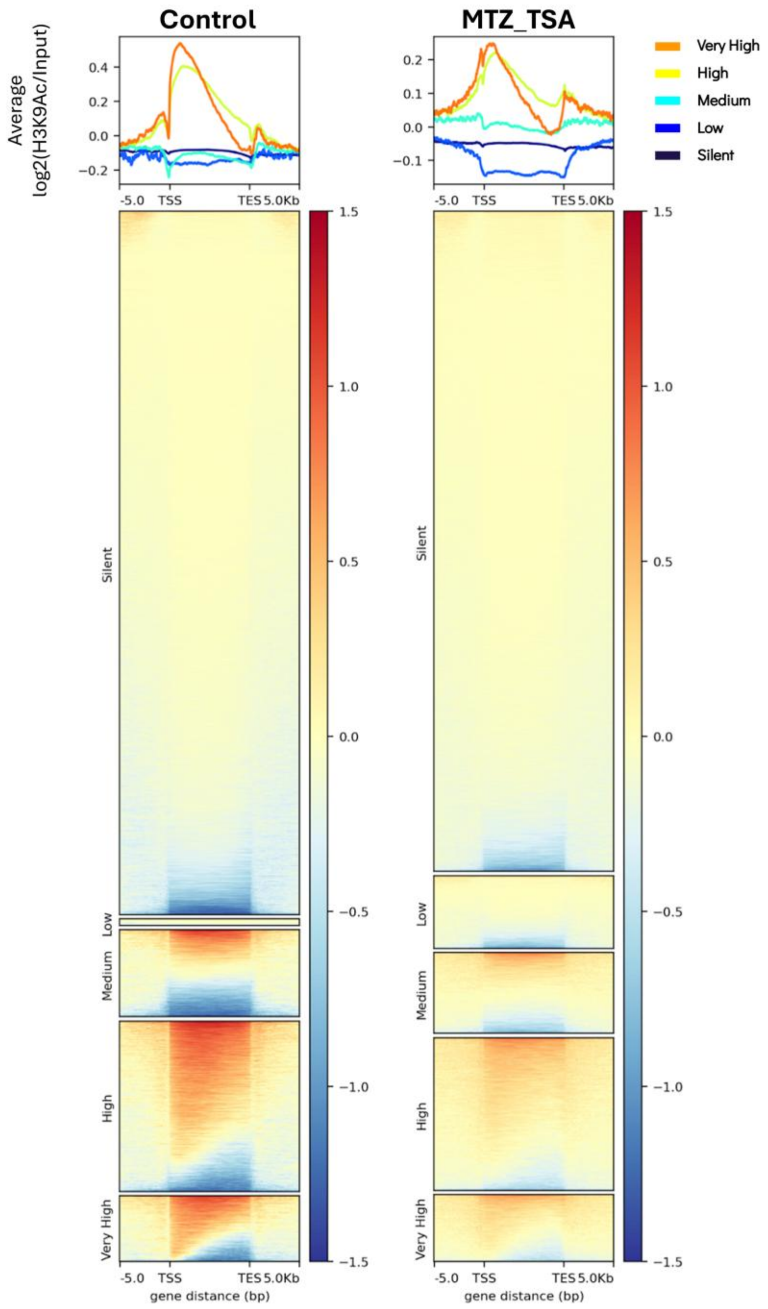

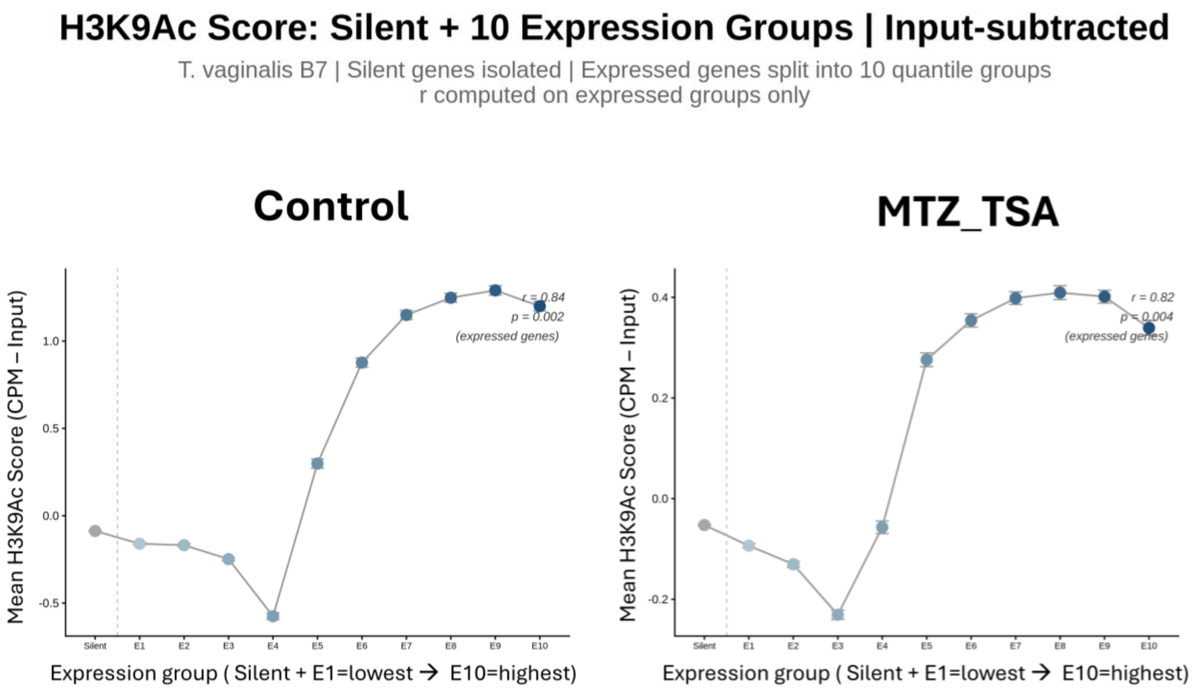

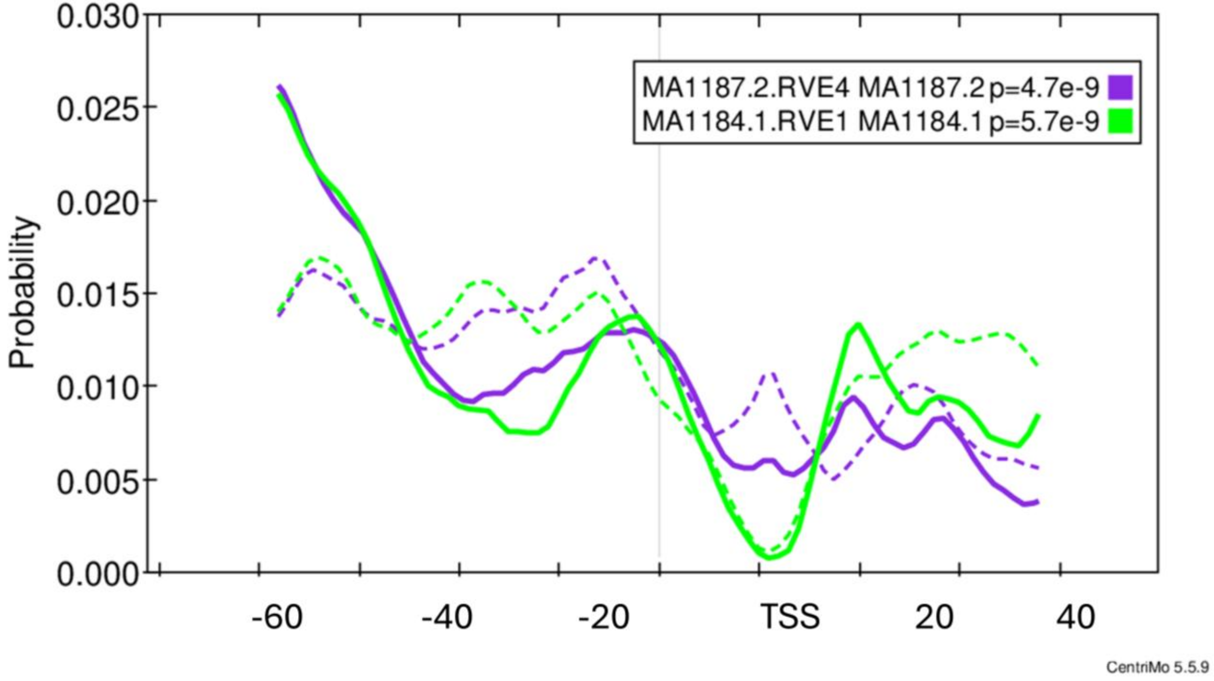
